# Nivolumab and Ipilimumab in Metastatic Melanoma Are Associated with Distinct Immune Landscape Changes and Response-Associated Immunophenotypes

**DOI:** 10.1101/2020.01.13.905430

**Authors:** DM Woods, AS Laino, A Winters, J Alexandre, V Rao, SS Adavani, JS Weber, PK Chattopadhyay

## Abstract

The reshaping of the immune landscape by nivolumab (NIVO) and ipilimumab (IPI) and its relation to patient outcomes is not well-described. We used high-parameter flow cytometry and a novel computational platform, CytoBrute, to define immunophenotypes of up to 15 markers to assess peripheral blood samples from metastatic melanoma patients receiving sequential NIVO>IPI or IPI>NIVO. The two treatments were associated with distinct immunophenotypic changes and had differing profiles associated with response. Only two immunophenotypes were shared but had opposing relationships to response/survival. To understand the impact of sequential treatment on response/survival, phenotypes that changed after the initial treatment and differentiated response in the other cohort were identified. Immunophenotypic changes occurring post-NIVO were predominately associated with response to IPI>NIVO, but changes occurring post-IPI were predominately associated with progression with NIVO>IPI. Among these changes, CD4+CD38+CD39+CD127-GARP- T-cell subsets were increased after IPI treatment and were negatively associated with response/survival for the NIVO>IPI cohort.

## Introduction

Immunotherapies have had remarkable success in treating a variety of malignancies including metastatic melanoma. However, many patients do not benefit from these treatments, and mechanisms differentiating patient outcomes remain elusive. The immune checkpoint antibodies ipilimumab (IPI) and nivolumab (NIVO) block ligation of the CTLA4 and PD1 inhibitory receptors, respectively, expressed on T-cells. Consequently, proper T-cell receptor (TCR) signaling and acquisition of effector function is restored to tumor-reactive T-cells^1^. These agents are given systemically, so they also exert effects on other, non-tumor-reactive T-cell populations, likely altering the immune landscape. However, the impact of these agents on the immune landscape and its potential relation to patient outcomes remains under investigation.

Combination therapy with IPI and NIVO results in a 55% response rate and 52% overall survival at five years, superior to either single agent, albeit at the cost of increased immune-related toxicity^2, 3^. To reduce treatment-related toxicity, the sequential administration of NIVO and IPI for the treatment of metastatic melanoma was evaluated in the Checkmate CA209-064 trial^4^. In this trial, significant differences in patient outcomes were observed; patients receiving IPI for 12 weeks, followed by NIVO for 12 weeks (IPI>NIVO) with subsequent NIVO maintenance had significantly lower response rates and overall survival compared to the NIVO>IPI sequence.

Defining biomarkers of patient outcome for immunotherapy has been the focus of much recent effort. Several biomarkers differentiating patient response to checkpoint inhibition have been described including tumor and infiltrating cell PDL-1 expression, tumor mutation burden, expression of a tumor gamma interferon-induced gene expression signature, increased levels of tumor infiltrating CD8+ T lymphocytes, and others^5^. Interestingly, there are few biomarkers associated with response to both αPD1 and αCTLA4. For some biomarkers, such as TCR diversity, T-cell memory subsets and frequency of circulating Tregs, there is a reciprocal association with response for the two therapies^6,7,8,9^. While useful for identifying rational targets for combination therapy, these biomarkers have failed to stratify patient responses well enough to have clinical utility for prediction of outcome. To accomplish this, multiple biomarkers that capture the complexity of individual immune responses may be required. Emerging high-dimension, single-cell technologies offer the ability to probe the immune response at an unprecedented depth and define novel biomarkers of outcome.

High-parameter, single-cell technologies are powerful tools to address whether variations in the immune landscape may be associated with clinical benefit. Their utility in biomarker discovery has been highlighted by recent publications detailing signatures and complex immunophenotypes associated with metastatic melanoma patient outcomes. For example, utilizing single-cell RNA-Seq, tumor transcriptional signatures associated with immunological exclusion and CD8+ T-cell transcriptional states associated with checkpoint immunotherapy resistance have been shown^10^. Mass cytometry, another high-dimensional, single cell technology, has also shown promise in discovering immune cell phenotypes associated with immunotherapy response/resistance. Using this technology, the frequency of monocyte populations has been shown to be associated with patient response to immunotherapy and overall survival^11^.

For the work presented here, we utilized a new, rapid computing platform for enumerating complex cell populations in single cell datasets, described by combinatorial expression of up to 15 markers. Using this approach, we describe the effects of NIVO and IPI on the peripheral blood immune landscape of metastatic melanoma patients and determine the association of those effects with response to therapy and overall survival. We used high-parameter (22- to 27-color) flow cytometry to assess peripheral blood specimens from patients treated with sequential NIVO>IPI or IPI>NIVO. By using paired baseline and week 13 specimens, we were able to directly compare the immunophenotypic changes accompanying treatment with single agent NIVO or IPI. We utilized a novel computational platform, CytoBrute, to assess frequencies of cell types defined by combinatorics for CD4+ and CD8+ T-cells, and myeloid cells. This methodology allowed us to identify clusters of immunophenotypically-related cells that were associated with each treatment and with patient outcomes.

## Materials & Methods

### Patient samples and processing

Cryopreserved peripheral blood mononuclear cells (PBMC), obtained from samples collected at baseline, and week 13 of the CA-209-064 clinical trial (ClinicalTrials.gov identifier NCT01783938), were thawed, washed, and stained for flow cytometry in a single batch. Viability for all samples was >85%. Samples were fixed in 0.5% paraformaldehyde. Baseline and week 13 PBMC obtained from patients treated with nivolumab monotherapy as part of the CA209-006 (ClinicalTrials.gov identifier NCT01176461) were similarly assessed in validation experiments.

### Flow cytometry

Four, 24+ parameter antibody panels (Supplemental Table 1) were developed using ColorWheel software, an automated flow cytometry panel design tool that proposes candidate panels based on spillover-spreading error, dye brightness, and antigen density. Data were collected on a custom BD FACSymphony A5 30-parameter flow cytometer, using a 96-well plate reader.

### Data analysis

After compensation for overlap in fluorescent spectra, removal of events representing fluorescence aggregates, and exclusion of cell doublets and dead cells, we identified fluorescence intensity thresholds (i.e., gates) that distinguished positive from negative expression of each marker studied. Fifteen markers were chosen from each panel for further analysis, based on the strength of antibody staining, variability across donors, and biological interest. CytoBrute (RocketML, Portland, OR), an adaptation of the R-algorithm FlowType based on RocketML’s rapid computing technology, was then used to measure the number of cells expressing every individual marker and every combination of 2 through 15 markers within the datafile. CytoBrute reported the top 1000 most frequent combinatorial phenotypes in each data file, along with those found amongst the top 1000 in some patients but not the others. In total, approximately 80,000 cell populations were compared across patient groups and time points.

Many of these populations were not independent (i.e., one population shared markers with another); therefore, we developed approaches to better characterize and summarize the results of group-wise comparisons. We clustered cell populations using the umap R package with default settings to group similar phenotypes^12^. Specifically, we determined cluster number by calculating a consensus from all non-graphical methods included in the Nbclust R package^13^, and then employed an unsupervised clustering algorithm (k-means) to identify clusters.

### Statistical Analysis

To compare the longitudinal immunologic effects of IPI and NIVO, univariate p-values for each measured immune feature were calculated from Week 0 and week 13 samples, within each treatment regimen, using a paired Wilcoxon test. Similarly, we identified associations between immunophenotypes and treatment response by grouping patients as responders or progressors using RECIST 1.1 criteria as reported for the Checkmate 064 trial^4^ and performing Mann-Whitney U tests comparing the frequencies of phenotypes across groups. Kaplan-Meier survival curves were also generated to compare patients with high (above the median) and low (below the median) frequencies of immunophenotypes of interest. Survival curves were compared using Mantel-Cox test and hazard ratios (HR) determined using a Mantel-Haenszel test.

Common methodologies for multiple comparison adjustments are not appropriate for our dataset, because the phenotypes generated by CytoBrute are not independent. Moreover, non-parametric statistical testing was utilized, and, as such, the lowest achievable p-values would not reach significance with traditional multiple comparison adjustments. Therefore, we used the above statistical tests to screen for populations differing across study groups at the p<0.05 level, and then confirmed results for select populations using a validation cohort of patients and/or by feeding populations into elastic net (EN) regularized regression models^14^ with leave-one-out (LOO) cross validation. EN regularized regression was performed with the R package glmnet^15^. The EN algorithm was chosen for its ability to handle a feature set with high collinearity. For a given EN model, the cv.glmnet function was used to obtain a lambda value within one standard error of the minimum mean cross-validated error in order to avoid overfitting. To robustly identify the altered immune features, we repeated this process a total of 100 times. For each patient, we then averaged model predictions for every model of the 100 in which they were part of the test set to get a final blinded prediction for each patient. We used this final prediction to calculate a receiver operator characteristic (ROC) and area under the curve (AUC) using the pROC R package^16^. Finally, we recorded the total number of times a given immune feature was included in all 100 models to get a frequency of selection, a proxy for the importance or predictive power of that feature.

## Results

### High dimension flow cytometry and combinatorics characterize immunophenotypes in metastatic melanoma patients

In the Checkmate 064 trial, metastatic melanoma patients received treatment with sequential NIVO>IPI or the reverse sequence IPI>NIVO (**Figure 1A**). To determine how NIVO and IPI shaped the immune landscape, we performed high-dimension flow cytometry on baseline and week 13 PBMC samples (i.e., after the first drug in each sequence). The markers assessed in four separate staining panels are shown in **Supplemental Table 1**.

**Figure 1.**
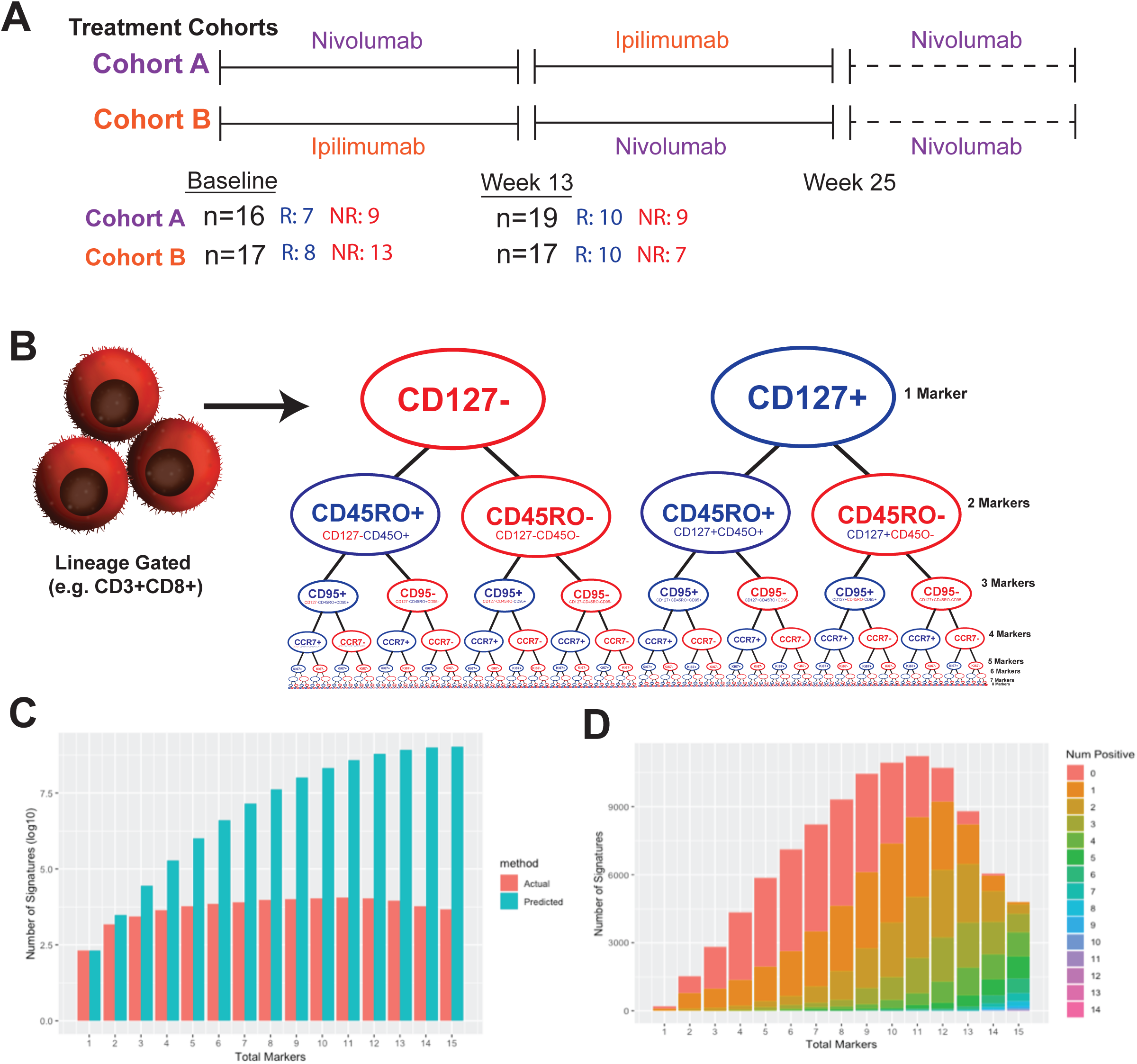
Overview of Approach. **(A)** PMBC samples were obtained from metastatic melanoma patients treated as part of the CA209-064 clinical trial. Patients were treated with sequential NIVO>IPI (Cohort A) or IPI>NIVO (Cohort B). Samples were collected prior to treatment (baseline) and post-first agent (week 13). The number of samples for each cohort and timepoint, broken down by patient response (responders in blue; non-responders in red), are given. **(B)** A generalized illustration of how the data were assessed using combinatorics and CytoBrute is shown. Briefly, for each cell population (e.g., CD3+CD8+ in the example illustration), the proportion of cells expressing each markers, and all combinations up to 15 markers, was assessed. This was performed for four separate flow cytometry panels. **(C)** The number of theoretical immunophenotypes for 1 to 15 markers in complexity is shown by blue bars and the corresponding number of actual immunophenotypes measured in the data set are shown in red. **(D)** For the actual immunophenotypes measured, the number of positive markers measured at each increment of complexity is shown. As per the legend to the right of the graph, immunophenotypes containing no positive markers are shown in red, those with one in orange and so on.

We first performed tSNE analyses (**Supplemental Figure 1**) to compare treatment regimens and time points. While differences were noted between responders and non-responders and between week 0 and week 13, the significance of the changes reported by the tSNE analysis was difficult to ascertain, particularly in identifying treatment induced changes given the non-paired nature. It was also difficult to exhaustively identify all the changes depicted in a tSNE graphic using only visual approaches. We also noted that, for the highly heterogeneous and diverse cell populations studied in our flow cytometry panels, tSNE fails to produce clearly separated “islands” of cells, further complicating the identification of cell types that differ by patient group or time point. These analyses demonstrate the difficulty and subjectivity of identifying changes in cell frequencies by tSNE.

To evaluate immunophenotypes in a more comprehensive manner, we used CytoBrute, a novel computational approach designed for analysis of high-parameter flow data sets. Briefly, each flow cytometry data file was gated on immune cell lineages (i.e. viable CD3+CD4+, CD3+CD8+, and leukocytes) of interest. Data were then analyzed using CytoBrute, which creates Boolean combinations for all possible phenotypes up to fifteen markers and calculates the frequency of cells expressing each combination of markers. **Figure 1B** illustrates the basis of this approach. Although this approach can theoretically generate up to fourteen million phenotypes (3^15^ subsets, considering positive, negative and neutral expression), many of these cell types are exceedingly rare or non-existent, so we selected the top 1,000 most frequent phenotypes per sample, for each antibody panel. As each sample had different immunophenotypes comprising the top 1,000, non-overlapping frequencies were also calculated, resulting in ∼100,000 immunophenotypes evaluated per antibody panel. The frequencies of immunophenotypes observed each panel are depicted as mosaic plots in **Supplemental Figure 2**. Each tile is sized to indicate the proportion of cells expressing a particular phenotype, and tiles are sorted from most frequent phenotypes (upper left corner) to least frequent (lower right corner). The tiles are color-coded by the number of markers defining the phenotype. The mosaic plot demonstrates the relevance of phenotypes defined by a high number of markers (i.e. phenotypes defined by eight to 15 markers, green and red boxes, most easily identified for Panel 1), which are interspersed amongst the most frequently observed phenotypes. Notably, there is a small proportion of phenotypes that are rarely observed or absent (gray boxes, lower right corner), demonstrating that the majority of potential immunophenotypes, both frequent and infrequent, are represented within the data.

### Changes in the peripheral blood immune landscape following NIVO or IPI treatment

We assessed the effects of NIVO and IPI on peripheral blood immune cells derived from metastatic melanoma patients by assessing baseline and week 13 paired samples using Wilcoxon signed-rank tests. With NIVO treatment, 1,744 immunophenotypes increased from baseline (pre-treatment), while 2,284 were decreased at p≤0.05 (**Figure 2A**). To dissect how the altered immunophenotypes were related, we projected significantly changed immunophenotypes into a two-dimensional Uniform Manifold Approximation (UMAP), and then clustered the data using k-means clustering. As shown in **Supplemental Figure 3A**, eight clusters were identified amongst the immunophenotypes increased post-NIVO. The frequency for the top 15 markers represented in each of these clusters is shown in **Supplemental Figure 3B**. Orange bars represent markers that are expressed (+); gray bars represent markers that are not expressed (-). The length of the bar denotes the percentage of immunophenotypes in that cluster expressing the corresponding marker. For example, the most common marker in the immunophenotypes composing cluster 6 is CD38+, expressed by nearly all of the immunophenotypes in that cluster. The second most common marker comprising this cluster is GITR-. Both CD4+ and CD8+ T-cells are represented in this cluster. Cell types that increased after NIVO include: T-cells expressing the ectonucleotidases CD38 and CD39 (clusters 1, 6, and 7) and CD73 (cluster 1), T-cells with a naïve-like phenotype (cluster 2), as well as cell types that are not well-defined by the markers we measured (clusters 3-5 and 8), Eight clusters were also identified for the immunophenotypes decreasing post-NIVO, as shown in **Supplemental Figure 3C**. The phenotypic composition of these clusters is shown **in Supplemental Figure 3D**. Cell types that decrease with NIVO were predominately characterized expression of markers associated with central memory T-cells (e.g. CD45RO+, CCR7+, CD127+, CD95+).

**Figure 2.**
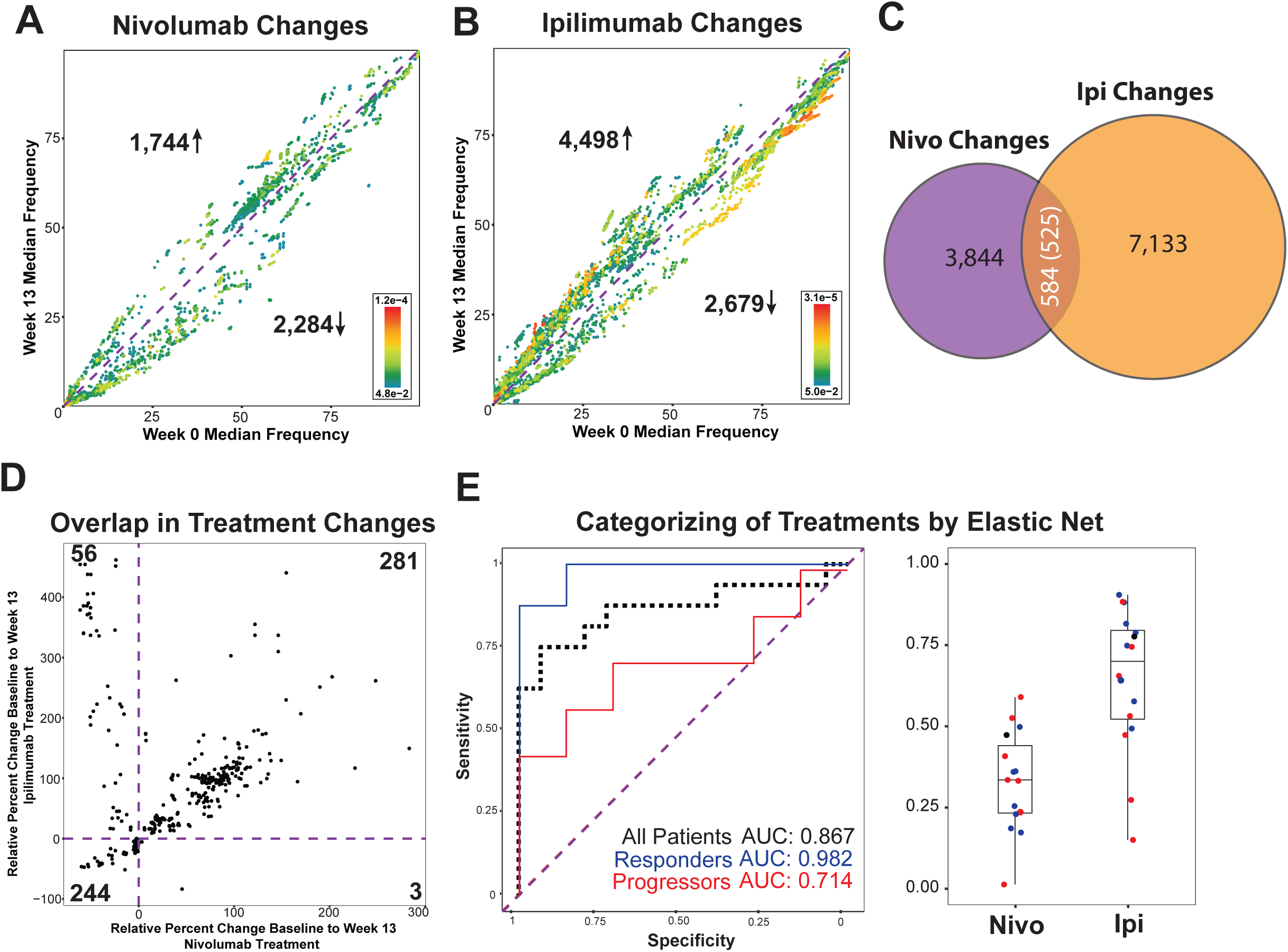
Nivolumab and ipilimumab differentially impact on peripheral blood immunophenotypes. **(A)** The median frequency at baseline (x-axis) and the week 13 median frequency (y-axis) are shown for immunophenotypes significantly elevated or diminished (p<0.05, Wilcoxon signed-rank test) in nivolumab-treated patients. Each dot represents an immunophenotype and is colored by p-value. The purple dotted line with a slope of one corresponds to no change in median frequency between baseline and week 13. **(B)** Ipilimumab-treated patient samples are shown, using a similar scheme. **(C)** Venn Diagram reporting the number of significantly changed immunophenotypes in each group and the overlap between them. The (525) immunophenotypes are those overlapping with changes in the same direction in both NIVO- and IPI-treated patient samples. **(D)** The median relative change from baseline to week 13 for nivolumab-treated patient samples (x-axis) and the relative change in ipilimumab-treated patient samples (y-axis) is shown for the 584 overlapping immunophenotypes. The purple dotted lines correspond to no change in median frequency. **(E)** The delta values (week 13 minus baseline) of the 584 overlapping phenotypes were used in an elastic net regularized regression model to categorize whether a paired patient sample received nivolumab or ipilimumab treatment. The receiver operator characteristic (ROC) and resulting area under the curve (AUC) for all paired samples is shown by the dotted black line in the left panel. The ROC and AUC for responding patient samples is shown in blue and for progressing patient samples in red. The model values for nivolumab and ipilimumab-treated paired patient samples are plotted in the right panel.

In IPI-treated patients, 4,498 immunophenotypes were increased and 2,679 were reduced relative to baseline with a p-value ≤0.05 (**Figure 2B**). For immunophenotypes increasing post-IPI, eight clusters were identified (**Supplemental Figure 4A**); these were defined by the markers listed in **Supplemental Figure 4B**. In contrast to NIVO-associated changes, many of these phenotypes were characterized by CD45RO+ and CD95+. Nine clusters were identified from the immunophenotypes decreasing post-IPI (**Supplemental Figure 4C**), for which the phenotypic breakdown is shown in **Supplemental Figure 4D**.

### NIVO and IPI have distinct impacts on the peripheral immunophenotypic landscape

To better compare the immunophenotypic effects of the two drugs, we examined which changes were common to both drugs versus unique to only one drug. For the two treatments, 584 immunophenotypes that changed overlapped (∼5%), as shown in **Figure 2C. Figure 2D** shows that of the 584 immunophenotypic changes common across drugs, 281 increased and 244 decreased. However, 56 immunophenotypes had reciprocal changes; these cell populations were expanded post-IPI but contracted post-NIVO. Only three phenotypes were increased post-NIVO but decreased post-IPI. **Supplemental Figure 5A** shows the immunophenotypes that increased with both treatments fell into seven clusters. A CD4+CD38+ T-cell phenotype predominated in clusters 2, 3, 4, and 7 (**Supplemental Figure 5B**). The 244 immunophenotypes that decreased with IPI or NIVO grouped into seven clusters (**Supplemental Figure 5C**), which included almost exclusively CD4+ immunophenotypes with the exception of cluster 7 (**Supplemental Figure 5D**). Six clusters represented the immunophenotypes decreasing post-NIVO but increasing post-IPI (**Supplemental Figure 5E**). Clusters 1 and 4 were composed of CD4+OX40+Ki67+ T-cells (**Supplemental Figure 5F**).

We next evaluated whether changes in circulating immunophenotypes could distinguish treatment regimens. To do so, the difference in frequency between week 13 and week 0 (delta value) was calculated for each immunophenotype. These delta values were then fed into an EN regularized regression model with repeated cross validation. Based on paired changes of the 584 identified overlapping phenotypes from all patient samples, the model was able to predict whether paired patient samples were from those who received NIVO or IPI with an area under the curve (AUC) of 0.867. The corresponding receiver operating characteristic (ROC) is shown by the black dotted line in the left panel of **Figure 2E**. The model values for individual paired samples are shown in the right panel of **Figure 2E**. Patient outcomes were added as a color dimension with responding patients (partial or complete response according to RECIST 1.1 criteria) in blue and progressing patients in red. Using the EN model determined by all patient samples, we then determined the ROC for responding and progressing patients separately. As shown in the left panel of **Figure 2E**, an AUC of 0.982 was achieved in responding patients (blue line) and an AUC of 0.714 in progressing patients, suggesting that patients who respond to therapy have more distinct immune changes than non-responders.

### Peripheral blood immunophenotypes at baseline and post-treatment are associated with patient outcomes after NIVO>IPI sequential therapy

We next sought to determine if baseline and/or week 13 (post-first agent) peripheral blood immunophenotypes were associated with patient outcomes. In baseline samples from NIVO>IPI treated patients (left panel, **Figure 3A**), 260 signatures were associated with both response to therapy (p≤0.05, Mann-Whitney U-test comparing responders and progressors) and overall survival (divided above and below median frequency, p≤0.05, Mantel-Cox test). Each dot represents a significant immunophenotype and is colored by the associated p-value from the comparison of frequency differences between responders and progressors. The x-coordinate is the median frequency of the immunophenotype in progressors and the y-coordinate is the corresponding median frequency in responders. The 61 immunophenotypes significantly elevated at baseline in responding patients formed five clusters, as shown in **Supplemental Figure 6A**. Cluster 1 consisted of CD8+CD95+PD1-CD25-T-cells, Clusters 2 and 4 consisted of CD4+CD45RA+CD127+HELIOS-CD73-CD49B-CD38-T-cells, and Cluster 5 contained CD4+ or CD8+ T-cells expressing LAG3 (**Supplemental Figure 6B**). The 199 immunophenotypes that were lower in responding patients (and therefore higher in those progressing patients) formed eight clusters as shown in **Supplemental Figure 6C**. Clusters 1, 3 and 7 were composed of CD4+CD38+CD39+CD127-T-cells (**Supplemental Figure 6D**). Clusters 2, 4, 5, 6 and 8 were composed of CD95+ expressing CD4+ T-cells.

**Figure 3.**
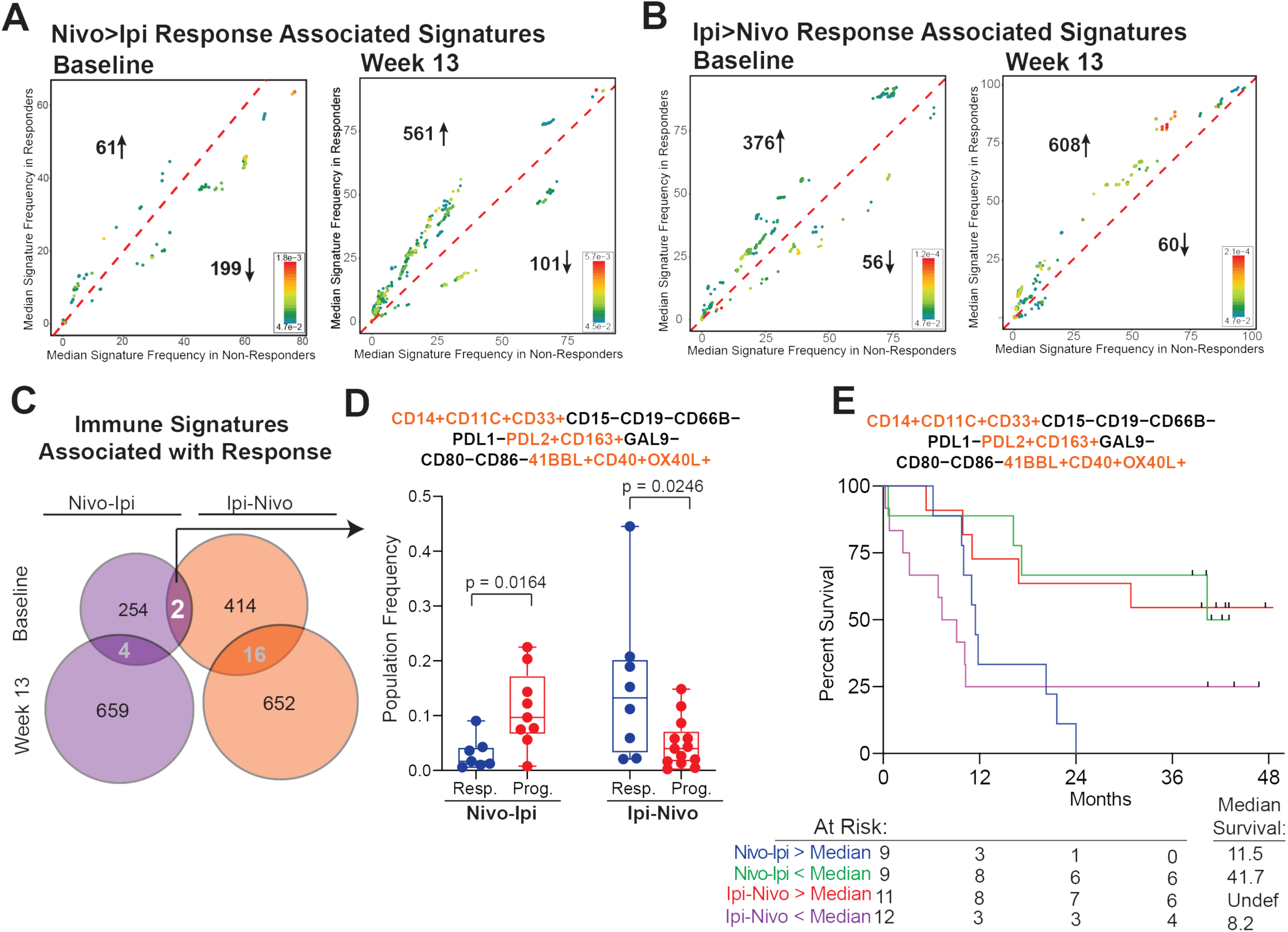
Immunophenotypes associated with patient response are distinct in nivolumab and ipilimumab sequentially treated patients. **(A)** The median frequency of immunophenotypes that are significantly different for both response and overall survival in non-responding patients are shown on the x-axis and in responding patients on the y-axis for nivolumab>ipilimumab treated patient samples. Each dot represents an immunophenotype and is colored by significance differentiating response. The purple dotted line with a slope of one corresponds to no change in median frequency. Significantly different immunophenotypes in baseline patient samples are shown in the left panel and significantly different immunophenotypes in week 13 patient samples are shown in the right panel. **(B)** Immunophenotypes for ipilimumab>nivolumab treated patients are likewise shown. **(C)** A Venn Diagram is shown with the number of immunophenotypes significantly different in each cohort and time point. **(D)** A graph of one of the two related significantly different immunophenotypes overlapping between nivolumab>ipilimumab and ipilimumab>nivolumab treated patients at baseline is shown. Frequencies of the populations shown are plotted by cohort and response. Each dot represents an individual patient sample. **(E)** A survival plot for this immunophenotype is also shown. Patients were stratified based on median frequency of the immunophenotype. Nivolumab>ipilimumab treated patients with greater than median frequencies are shown in blue and less than median frequency in green. Ipilimumab>nivolumab treated patients with greater than median frequencies are shown in red and less than median are shown in purple.

Post-NIVO (week 13), 662 immunophenotypes were found to be associated with patient outcomes in those receiving sequential NIVO>IPI treatment (right panel, **Figure 3A**). In contrast to baseline response-associated immunophenotypes which were predominately CD4+, nearly all the post-NIVO response-associated immunophenotypes were CD8+, in particular those found at higher frequencies in responders. Amongst these immunophenotypes, 561 were increased in frequency in responding relative to progressing patients. As shown in **Supplemental Figure 6E**, ten clusters were formed from these immunophenotypes. Combinations of CCR7, CD127, CD45RO and CD95 positive CD8+ T-cells comprised Clusters 2, 3, 4, 5, 6, 8 and 10. A total of 101 immunophenotypes forming seven clusters were found to be decreased in responding relative to progressing patients (**Figure 3A** right panel and **Supplemental Figure 6G**). As shown in **Supplemental Figure 6H**, Clusters 1, 2, and 6 were composed of CD8+CD45RA+ T-cells. Collectively, a number of immunophenotypes that were associated with patient response and survival and were grouped into several immunophenotypically-related clusters were identified. These included elevated levels of naïve-like (CD45RA+, CD127+) T-cells and CD8+LAG3+ phenotypes at baseline and elevated levels of central-memory-like (CD45RO+, CCR7+, CD127+, CD95+) T-cells post-NIVO. A population of CD4+CD38+CD39+ T-cells at baseline was also associated with progression and shorter survival.

### Peripheral blood immunophenotypes at baseline and post-treatment are associated with patient outcomes after IPI>NIVO therapy

In IPI>NIVO treated patients, 432 baseline immunophenotypes were associated with response to treatment and survival (**Figure 3B**, left panel). Of these, 376 were elevated in frequency in responding relative to progressing patients. These 376 immunophenotypes formed eight clusters and were a mixture of CD4+ and CD8+ T-cell populations (**Supplemental Figures 7A and 6B**). Clusters 5 and 6 were composed of CD4+ T-cells expressing LAG3+ and GARP+. 56 phenotypes, forming five clusters, were decreased in frequency in responding patients (**Supplemental Figure 7C and 7D**). All clusters contained a predominance of CD45RO+ and CD95+ immunophenotypes, while Clusters 2, 4 and 5 also contained CCR7-expressing immunophenotypes.

At week 13, after IPI treatment (**Figure 3B**, right panel), 668 immunophenotypes were associated with treatment response and survival, with 608 elevated and 60 at lower frequencies in responding compared to progressing patients (**Figure 3B**, right panel). As shown in **Supplemental Figure 7E**, immunophenotypes higher in relative frequency in responders formed seven clusters. CD4+ and CD8+ T-cell immunophenotypes with CCR7+ expression comprised Clusters 1, 2, 3, 5 and 7 (**Supplemental Figure 7F**). The 60 immunophenotypes with lower frequencies in responders formed 6 clusters (**Supplemental Figure 7G**). These clusters were mixed populations of CD4+ and CD8+ T-cells (**Supplemental Figure 7H**). Cluster 2 included T-cells expressing CD38+ and CD39+.

### Distinct immunophenotypes are associated with patient response in NIVO>IPI and IPI>NIVO treated patients

We next compared the immunophenotypes associated with outcome between NIVO>IPI and IPI>NIVO patients. As shown in **Figure 3C**, only two (related) immunophenotypes were associated with response in both cohorts. The associations were reciprocal between the two cohorts. These cells were CD14+CD11C+CD33+CD15-CD19-CD66B-PDL1-PDL2+CD163+GAL9-CD80-CD86-41BBL+CD40+OX40L+ and an identical immunophenotype in which CD66B was not measured. The frequency of this immunophenotype in responding and progressing patients for each treatment cohort is shown in **Figure 3D**. In NIVO>IPI treated patients, higher frequencies of these immunophenotypes were associated with progression (p=0.0164), while in IPI>NIVO treated patients, higher frequencies were associated with patient response (p=0.0246). This was reflected in the survival curves shown in **Figure 3E**. Frequencies of these phenotypes above the median were associated with shorter survival in NIVO>IPI treated patients (p=0.0050, HR 95% CI: 1.681-18.74) but with prolonged survival in IPI>NIVO treated patients (p=0.0463, HR 95% CI: 1.019-9.038).

### NIVO-associated immune landscape changes favor response-associated immunophenotypes in the IPI>NIVO cohort

In the Checkmate 064 trial, the NIVO>IPI treatment arm had greater rates of response and overall survival compared to the IPI>NIVO treatment arm^4^. We hypothesized that the immunophenotypic impact of treatment with NIVO or IPI may have altered the immune landscape in a manner that influenced subsequent response or resistance to the second agent. We tested this hypothesis by determining the overlap between the immunophenotypes that changed significantly after the first treatment in the regimen and those associated with treatment response and survival in the opposing cohort. As depicted in **Figure 4A**, of the 3,959 cell populations that were significantly altered after NIVO, four overlapped with those that were associated at baseline with response and survival in IPI>NIVO treated patients. However, none of the markers we measured were expressed by the cells in these four immunophenotypes (i.e. no positive markers) (data not shown). An additional 95 overlapping immunophenotypes associated with treatment response/patient survival at week 13 in the IPI>NIVO cohort and changed after NIVO treatment were found. Ninety-nine percent (94/95) of these overlapping immunophenotypes were positively associated with response, with almost all (93/94) immunophenotypes increasing post-NIVO and were elevated at week 13 in responders to IPI>NIVO.

**Figure 4.**
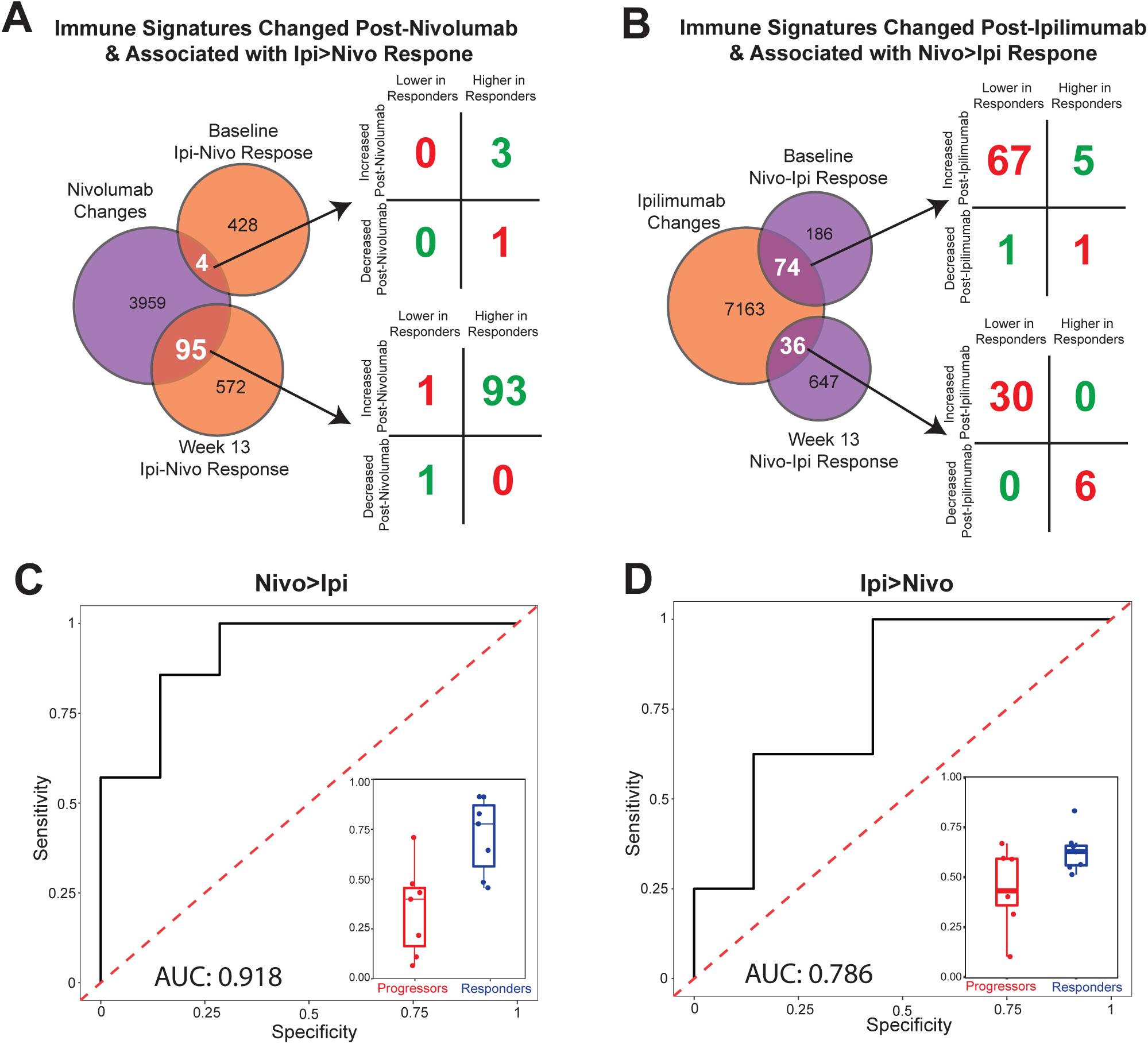
Ipilimumab-induced immunophenotypic changes are associated with lack of response to sequential nivolumab-ipilimumab. **(A)** A Venn diagram depicts the number of immunophenotypes significantly changed post-nivolumab (purple circle) and immunophenotypes associated with survival and response in ipilimumab>nivolumab treated patients at baseline (top orange circle) and post-ipilimumab (bottom orange circle). The immunophenotypes that overlap between these groups (intersection of Venn diagrams) are highlighted and analyzed to determine 1) the direction of change and 2) the relative representation in responding patients. This analysis was performed for the highlighted phenotypes at baseline (top four-square diagram) and post-ipilimumab (bottom four-square diagram). These four-square diagrams are laid out as follows: the number of immunophenotypes significantly increased post-nivolumab and lower in ipilimumab-nivolumab responding patients are shown in the top left quadrants; the number of immunophenotypes significantly increased post-nivolumab and higher in IPI>NIVO responding patients are shown in the top right quadrants; the number of immunophenotypes significantly decreased post-nivolumab and lower in IPI>NIVO responding patients are shown in the bottom left quadrants; the number of immunophenotypes significantly decreased post-nivolumab and higher in IPI>NIVO responding patients are shown in the bottom right quadrants. The notable result here is the overwhelming number of nivolumab-induced changes that are beneficial to IPI>NIVO outcomes (green numbers), compared to the few nivolumab-induced changes that are associated with poor IPI>NIVO survival and response. **(B)** A similar Venn diagram and four-square diagram are shown for significant ipilimumab changes and NIVO>IPI response/survival associated immunophenotypes is shown. The notable result here is that ipilimumab-induced changes overwhelmingly are associated with poor NIVO>IPI outcomes (red). **(C, D)** Delta values (week 13 - week 0) for the 210 immunophenotypes significantly changed post-nivolumab or post-ipilimumab and associated with response/survival in IPI>NIVO or NIVO>IPI treated patients were used in an elastic net regularized regression model to categorize patient response. **(C)** The ROC and AUC for patients in the NIVO>IPI-treated cohort are shown. The model values for non-responding and responding patients are plotted in the bottom right inlay. **(D)** Results for IPI>NIVO treated patients are likewise shown.

We next evaluated the composition of the 95 immunophenotypes identified above associated with treatment response/patient survival at week 13 in the IPI>NIVO cohort (**Supplemental Figure 8A**). The 93 immunophenotypes that were increased at week 13 post-NIVO and associated at baseline with response in IPI>NIVO (green dots in Supplemental Figure 8A) are described in **Supplemental Figure 8B**. These cells were all CD4+ and predominately CD45RO- and CCR7+. A representative population of CD4+CD45RO-CCR7+ T-cells is shown in **Figure 5A**. The leftmost panel shows that most patients had an increase in the frequency of this population post-NIVO treatment (p=0.035). The second panel shows that higher frequencies of this population in IPI>NIVO treated patients at week 13 were associated with response (p=0.0046). The second to right panel shows that IPI>NIVO patients with greater than the median frequency of CD4+CD45RO-CCR7+ T-cells at week 13 have longer overall survival (p=0.019, HR 95% CI: 1.343-27.42). To validate this result, we evaluated changes in the frequency of this immunophenotype in a separate cohort of metastatic melanoma patients treated with nivolumab monotherapy (ClinicalTrials.gov identifier NCT01176461). Forty paired patient samples were assessed. The rightmost panel of Figure 5A shows that in this cohort of patients treated with NIVO, CD4+CD45RO-CCR7+ T-cells increased in frequency post-NIVO (p=0.046).

**Figure 5.**
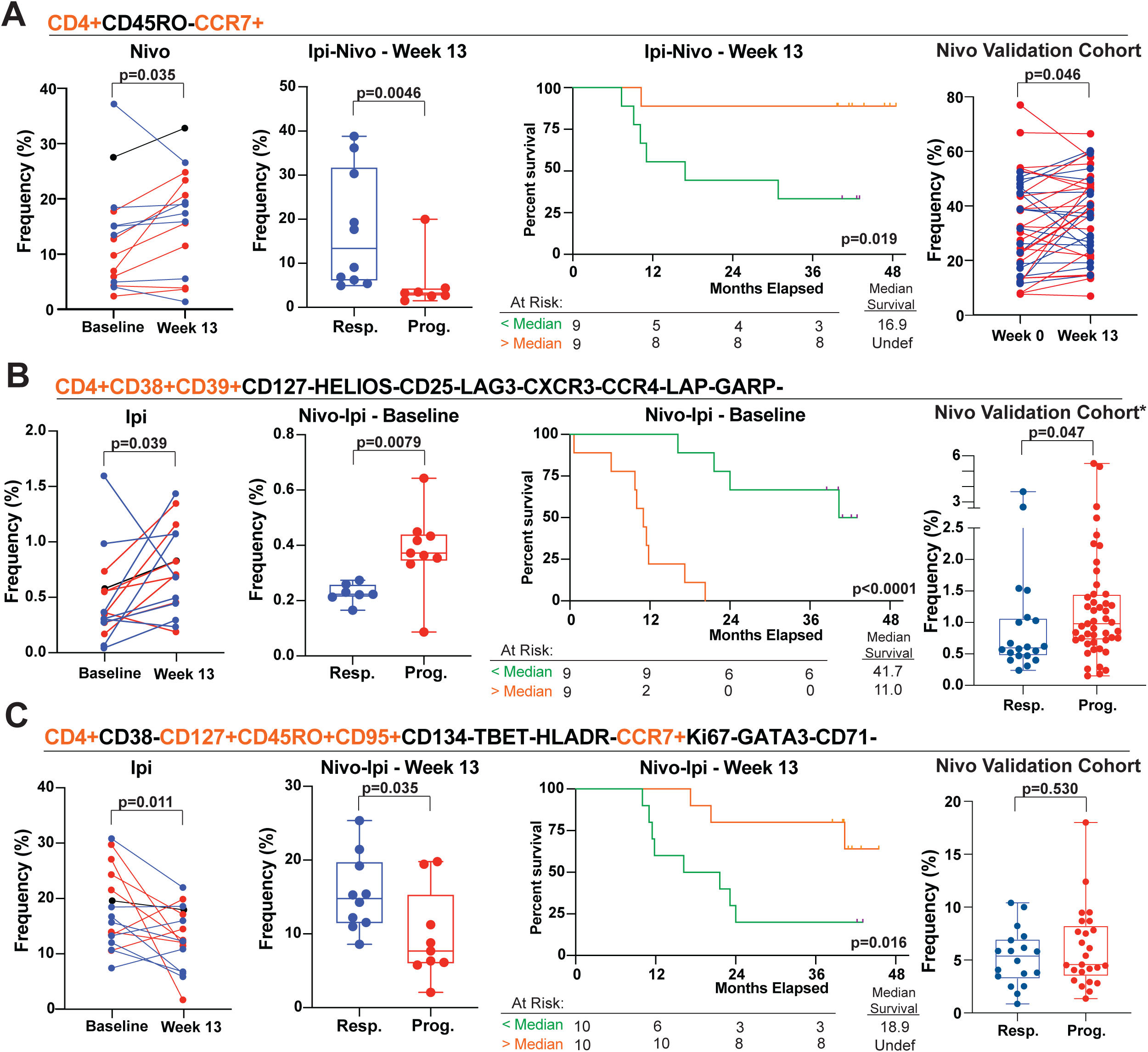
Ipilimumab-induced immunophenotypic changes are associated with lack of response to sequential NIVO>IPI patients. **(A)** A representative immunophenotype increased post-nivolumab and having an increased frequency in responding patients is shown. The leftmost graph shows the paired frequency changes in nivolumab-treated patients. Responding patients are represented by blue lines, non-responding patients with red lines, and non-evaluable patients in black. The second to left graph plots the week 13, post-ipilimumab frequencies of this population comparing responding and non-responding patients. The third panel is a survival plot for this immunophenotype. The rightmost graph shows the paired frequency changes in an independent validation cohort of nivolumab-treated patients. IPI>NIVO treated patients were stratified based on median frequency of the immunophenotype at week 13. Patients with greater than median frequencies are shown in orange and less than median frequency in green. **(B)** A representative immunophenotype increasing post-ipilimumab and having decreased frequency at baseline in NIVO>IPI responding patients is likewise shown. **(C)** A representative immunophenotype decreasing post-ipilimumab and having increased frequency at baseline in NIVO>IPI responding patients is likewise shown.

### IPI-associated immune landscape changes favor progression-related immunophenotypes in the NIVO>IPI cohort

In a similar assessment of the overlap between post-IPI changes and NIVO>IPI outcome-associated immunophenotypes, we identified 74 immunophenotypes that were altered by IPI treatment and whose frequency at baseline in the NIVO>IPI treatment arm was associated with response/survival (**Figure 4B**). Similarly, we identified 36 immunophenotypes that were altered by IPI, and whose presence at week 13 in the NIVO/IPI treatment arm was associated with response/survival (**Figure 4B**). **Supplemental Figure 8C** shows how these immunophenotypes cluster into groups of cell populations.

Unlike the NIVO-associated changes, IPI-associated changes had a negative association with NIVO response. At baseline, 92% of the overlapping phenotypes (68/74; **Figure 4B**) were associated with progression; 67 of these immunophenotypes were increased post-IPI and found at relatively lower frequencies in NIVO>IPI responding patients. As shown in **Supplemental Figure 8D**, these immunophenotypes were all CD4+CD38+CD39+GARP-CD127- and varied with respect to other markers. A representative population from these 67 immunophenotypes, CD4+CD38+CD39+CD127-HELIOS-CD25-LAG3-CXCR3-CCR4-LAP-GARP-T-cells, is shown in **Figure 5B**. The leftmost panel shows that this population was increased post-IPI (p=0.039). The second panel shows that significantly higher frequencies of this population are seen in progressing patients in the NIVO>IPI cohort (p=0.0079). The second to right panel shows that patients with greater than median frequency of this population also have significantly shorter survival (p<0.0001, HR 95% CI: 4.399-69.23). To independently validate the association of this immunophenotype with patient response, we assessed the frequency of CD4+CD38+CD39+CD127-GARP-T-cells in peripheral blood samples of metastatic melanoma patients treated with nivolumab monotherapy. Twenty responding and 47 progressing patient samples were assessed. Shown in the rightmost panel of **Figure 5B**, elevated levels of that immunophenotype were associated with progression of disease (p=0.047).

At week 13, all (36/36) of the phenotypes altered by IPI were associated with poor treatment response/survival for NIVO>IPI patients. Of the 36 immunophenotypes identified, IPI increased the frequency of 30 of these cell populations, but patients responding to NIVO>IPI had significantly decreased frequencies of these cells. **Supplemental Figure 8F** shows how these immunophenotypes clustered and **Supplemental Figure 8H** describes the marker composition of these clusters. The additional 6 overlapping immunophenotypes were decreased post-IPI but had elevated frequencies at week 13 in NIVO>IPI responding patients. These six immunophenotypes included both CD4+ and CD8+ T-cells and were predominately CD127+, CD95+, CD45RO+ and CCR7+. A representative phenotype from these six is shown in **Figure 5C**. The leftmost panel shows that this population was significantly decreased post-IPI (p=0.011). The middle panel shows that the frequency of this population at week 13 in NIVO>IPI patients is significantly higher in responders compared to progressors (p=0.035). The second to right panel shows the associated increased survival in patients with >median frequency of this population (p=0.016, HR 95% CI: 1.322-15.49). However, shown in the rightmost panel of Figure 5C, in a cohort of metastatic melanoma patients treated with NIVO, we were unable to confirm a relationship between reduced levels of these cells and disease progression (p=0.530).

### Changes in identified immunophenotypes correctly classify patient response in a cross-validated Elastic Net model

To determine if the changes in overlapping immunophenotypes described in Figure 4A and 4B were sufficient to predict patient outcomes, we used patient delta values for the 209 overlapping immunophenotypes and an EN model with LOO cross validation. As shown in **Figure 4C**, responding and progressing patients in the NIVO>IPI-treated cohort were accurately categorized with an AUC of 0.918. **Figure 4D** shows that patients in the IPI>NIVO treated cohort were correctly categorized as responders or progressors, but with lesser sensitivity and specificity, resulting in an AUC of 0.786. These results further support the importance of immunophenotypic changes resulting from treatment and their relation to patient response.

## Discussion

In the current study we utilized a novel and powerful approach to analyzing high-dimension flow cytometry data to assess the impact of the immune checkpoint inhibitors nivolumab and ipilimumab on the peripheral blood immune landscape. By assessing the frequencies of complex immunophenotypes *in lieu* of dimension-reduction analytical methods (e.g. tSNE), we were able to more precisely identify treatment-associated changes in the immunophenotypic landscape, identify response-associated immunophenotypes, and assess their relationships. Collectively, these data suggest that IPI and NIVO alter the peripheral blood immunophenotypic landscape of patients in distinct ways. Further, IPI-associated alterations overlapped with immunophenotypes associated with progression of disease and shorter survival in the NIVO>IPI cohort. Although the overlap occurred with the patient responses to sequential NIVO>IPI noted at week 25, overall responses deviated little from week 13 responses (post-NIVO alone)^4^. These results suggest that IPI-associated immune landscape changes may impair responses to subsequent NIVO and may explain the lower response rate and shorter survival seen in the IPI>NIVO cohort.

While the emergence of single-cell, high-dimension technologies has increased the ability to probe the anti-tumor immune response, approaches to analyze the complex data generated have failed to keep pace. Dimension-reduction techniques such as principal component analysis and tSNE are commonly used to visualize high parameter data, but these techniques do not report the specific combination of markers expressed by cell types. A critical, unanswered question in high-parameter data analysis is whether clustering and dimension-reduction algorithms sufficiently capture all the cell types that differ across study groups. Classically, to exhaustively examine cell phenotypes in lower parameter flow cytometry datasets, analysis has been performed by using combinatorics, a method that constructs all possible phenotypes from the markers measured. For example, using combinatorics, an *n*-color flow cytometry experiment would report the number of cells expressing each combination of expression (+) or lack of expression (-) for the *n* markers, resulting in 2*^n^* phenotypes. The use of a neutral condition for each marker allows assessment of shorter and simpler phenotypes, resulting in 3*^n^* phenotypes. The identification of simple phenotypes is critical for generating translatable discoveries from high parameter technology. After all, it would be unnecessarily complex and expensive to develop clinical tests to detect cell populations that are defined by much more than three markers, when simpler surrogates might readily be identified within a high parameter dataset.

While combinatorics offers a means to precisely identify and quantify all cell subsets in a sample, it is computationally intensive. Even for a 10-parameter experiment measuring one million cells, combinatoric analysis using the R-based flowType algorithm requires four hours; in comparison, the same analysis is completed by our CytoBrute platform within two seconds. The improvement in computational time is attributable to the unique and proprietary distributive computing approach CytoBrute uses, and this approach is adaptable to other R-based algorithms and applications, including machine learning-based analysis of high parameter datasets.

There are shortcomings in this study that impact the interpretation of the data. Rather than using traditional FDR approaches, we utilized a non-multiple comparison adjusted p-value of <0.05 as a determination of significance. This approach was taken for several reasons. First, the number of samples available for assessment were limited, and in turn the lower threshold of p-values obtainable was limited. Second, the combinatoric nature of the CytoBrute approach creates non-independent measurements which would be highly overcorrected if using traditional multiple comparison corrections. However, given 1) the use of leave one out cross validation in EN models, 2) the validation of important identified immunophenotypes in an independent cohort, and 3) our focus on demonstrating that the impact of IPI and NIVO are distinct and that the immune landscape changes induced by IPI are associated with lower response to NIVO and shorter survival in the IPI>NIVO cohort, the conclusions of this study are supported by the data. Many response-associated immunophenotypes not overlapping with treatment changes were also identified that have not yet been evaluated in a validation cohort. While beyond the scope of this study, these represent potentially important biomarkers and are the subject of future validation efforts.

Several thousand significant changes in peripheral blood immunophenotypes were observed after NIVO or IPI treatment, highlighting the systemic impact of these agents. These changes were largely distinct with only <5% overlap in significantly changed immunophenotypes. Taken with data presented showing a lack of overlap in outcome-associated immunophenotypes between the two cohorts/treatments and published literature^6,7,8,9^ it is increasingly clear that the systemic immune impact and the mechanisms by which CTLA4 and PD1 blockade function are distinct.

Of the few overlapping immunophenotypes, both treatments increased populations of CD4+CD38+CD39+CD127-GARP-T-cells, which were found to be associated with poor outcomes. This population was also associated with progression in an independent cohort of NIVO-treated patients, validating the importance of this phenotype in patient response. To the authors’ knowledge, this population of T-cells has not been previously described. CD39 is an ectonucleotidase that converts extracellular ATP into adenosine and has been the focus of many studies demonstrating its roles in generating an immunosuppressive resulting in its emergence as a potential therapeutic target^17,18,19,20,21,22^. CD38 also functions as an ectoenzyme, both as a hydrolase and converts NAD+ to cyclic ADP-ribose. It is being assessed as a target in combination with immunotherapy^23, 24^. In agreement with our data, CD38 expression was recently shown to be upregulated post-immunotherapy and was found to be associated with negative outcomes in a murine checkpoint inhibition model and in human patients^25^. Based on the current knowledge of the function of CD38 and CD39 and our observation that CD4+CD38+CD39+ immunophenotypic clusters are associated with poor patient outcomes, we hypothesize that this population is immunosuppressive. Investigations of mechanistic relationship to patient response, the function of this population and the efficacy of targeting it are warranted by the data presented in this study and are ongoing.

In addition to the low level of overlap of immunophenotypic changes, no immunophenotypes associated with patient outcomes were found to be shared between the treatment sequences, with the exception of two related immunophenotypes. This further highlights the distinct impact of the two therapies. The two overlapping immunophenotypes were reciprocally associated with patient outcomes in the two cohorts. Based on the expression of CD14, CD11b and CD33, these cells are likely of myeloid origin^26^ and expressed a mixed inflammatory (e.g. 41BBL+, CD40+)^27, 28^ and suppressive phenotype (e.g. PDL2+, CD80-, CD86-)^29,30,31^. While beyond the scope of the present study, this cell population will be further interrogated in functional assays to determine potential mechanisms by which it could be associated with the disparate outcomes in the two treatment cohorts.

T-cell phenotypes associated with memory subsets were consistently and significantly associated with patient outcomes in both treatment cohorts. In the NIVO>IPI cohort, responding patients had higher frequencies of T-cells with a naïve phenotype (CD4+CD45RA+CD127+) at baseline and central memory phenotypes (CD4/8+CD45RO+CD127+CD95+CCR7+) post-NIVO. Conversely, at baseline, progressing patients had more differentiated effector T-cells (CD4+CD45RO+CD95+) and higher frequencies of CD8+CD45RA+ T-cells post-NIVO. These data suggest that the formation of memory T-cells may be important for the efficacy of NIVO. In contrast, in the IPI>NIVO cohort, responding patients had increased frequencies of CD4/8+CD45RO+CD95+ and CD4+CD45RA+CD95+ T-cells at baseline and increased frequencies of CD4+CD45RO-CCR7+ T-cells post-IPI. Several other immunophenotypic clusters were found to be associated with patient response and survival including CD4+CD45RA+CD127+, CD4+CD45RO+CD95+ and CD8+LAG3+ T cell populations.

Collectively, these data suggest that the impact of IPI and NIVO on the immunophenotypic landscape of patients is distinct and that the impact of IPI may be associated with resistance to subsequent NIVO therapy, consistent with poor outcomes in the IPI>NIVO cohort of Checkmate-064. In further support of this interpretation, in clinical trials the response rates to NIVO in patients progressing after IPI are lower than those in IPI-naïve patients^32,33,34,35,36^. However, these response rates are compared across different studies and lower response rates to NIVO in IPI-refractory patients may result from selection of immunotherapy-resistant patients. Response rates in the NIVO>IPI cohort were similar to concurrent NIVO and IPI, suggesting that concomitant IPI does not negatively impact NIVO efficacy. Regardless, the data presented herein raise concerns about the negative impact of IPI treatment prior to NIVO and warrant consideration in patient treatment decisions. Future studies will investigate immunophenotypic changes occurring with concurrent treatment and associated patient outcomes. Further, studies will need to address whether the immune landscape changes resulting from IPI are normalized over time and the duration that takes to occur. Future investigations will also need to investigate the function of, and potential value as biomarkers of immune cell populations that are associated with treatment outcomes in this study.

## Acknowledgements

We extended our appreciation to Bristol-Myers Squibb, including Christine Horak and Megan Wind-Rotolo, for all of their assistance and feedback in this study.

## Authorship Contributions

- David M. Woods: conceived and conducted experiments, analyzed and interpreted data, prepared manuscript.
- Andressa S. Laino: conceived and conducted experiments, analyzed and interpreted data, prepared manuscript.
- Aidan Winters: conceived and conducted experiments, analyzed and interpreted data, prepared manuscript.
- Jason Alexandre: conducted experiments, edited manuscript.
- Vinay Rao: managed creation of data analysis algorithm/platform (Cytobrute)
- Santi Adavani: created and implemented data analysis algorithm/platform (Cytobrute)
- Jeffrey S. Weber: oversaw project, conceived experiments, interpreted data, edited manuscript.
- Pratip K. Chattopadhyay: oversaw project, conceived experiments, analyzed and interpreted data, prepared manuscript.

## Supplemental Figure Legends

**Supplemental Figure 1.**
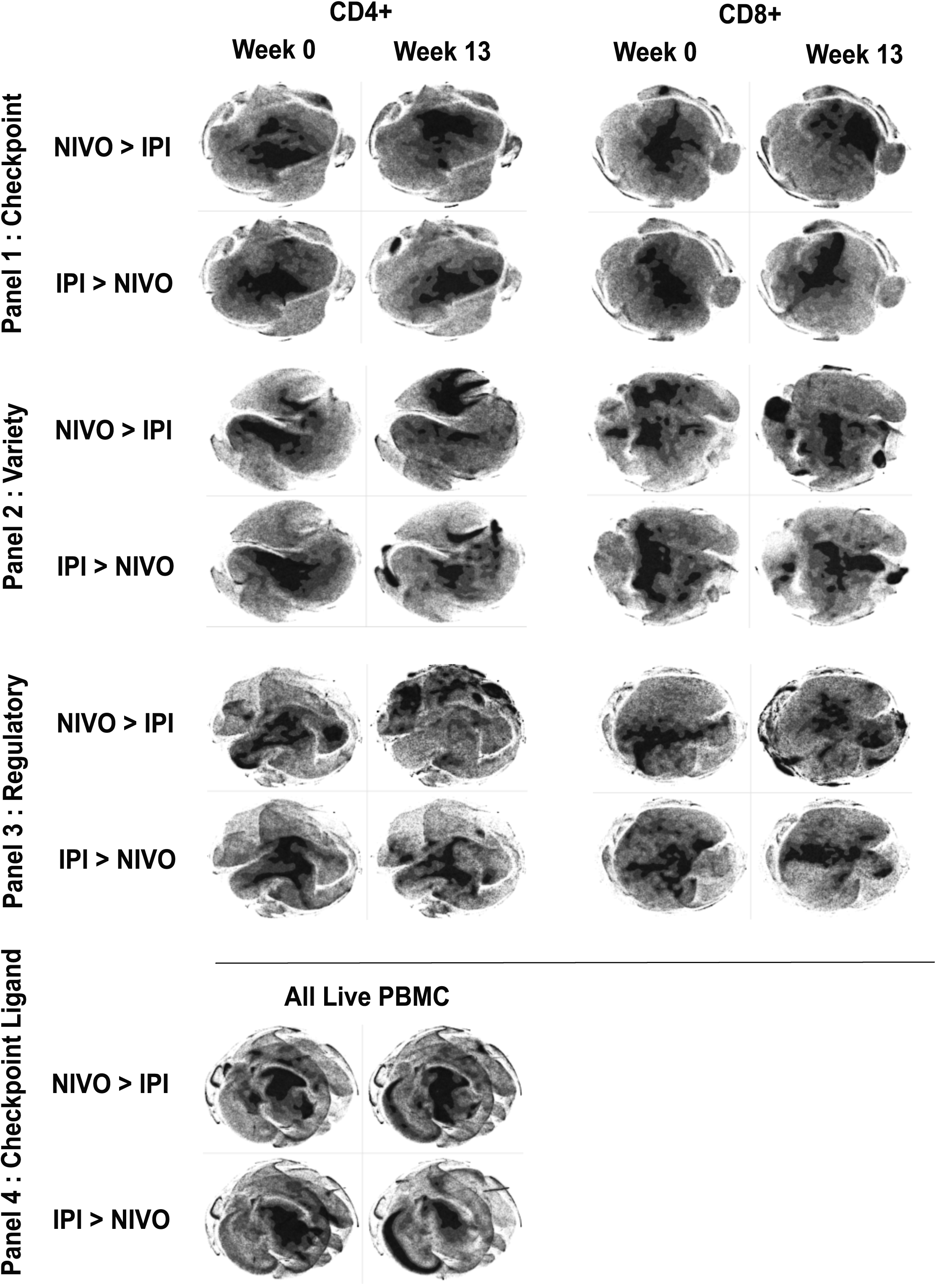
Tsne-based visualizations of immunophenotyping data from each antibody panel, for each patient group, for each time point. The difficulty in completely and comprehensively identifying all areas of the graphic that differ across study groups is evident.

**Supplemental Figure 2.**
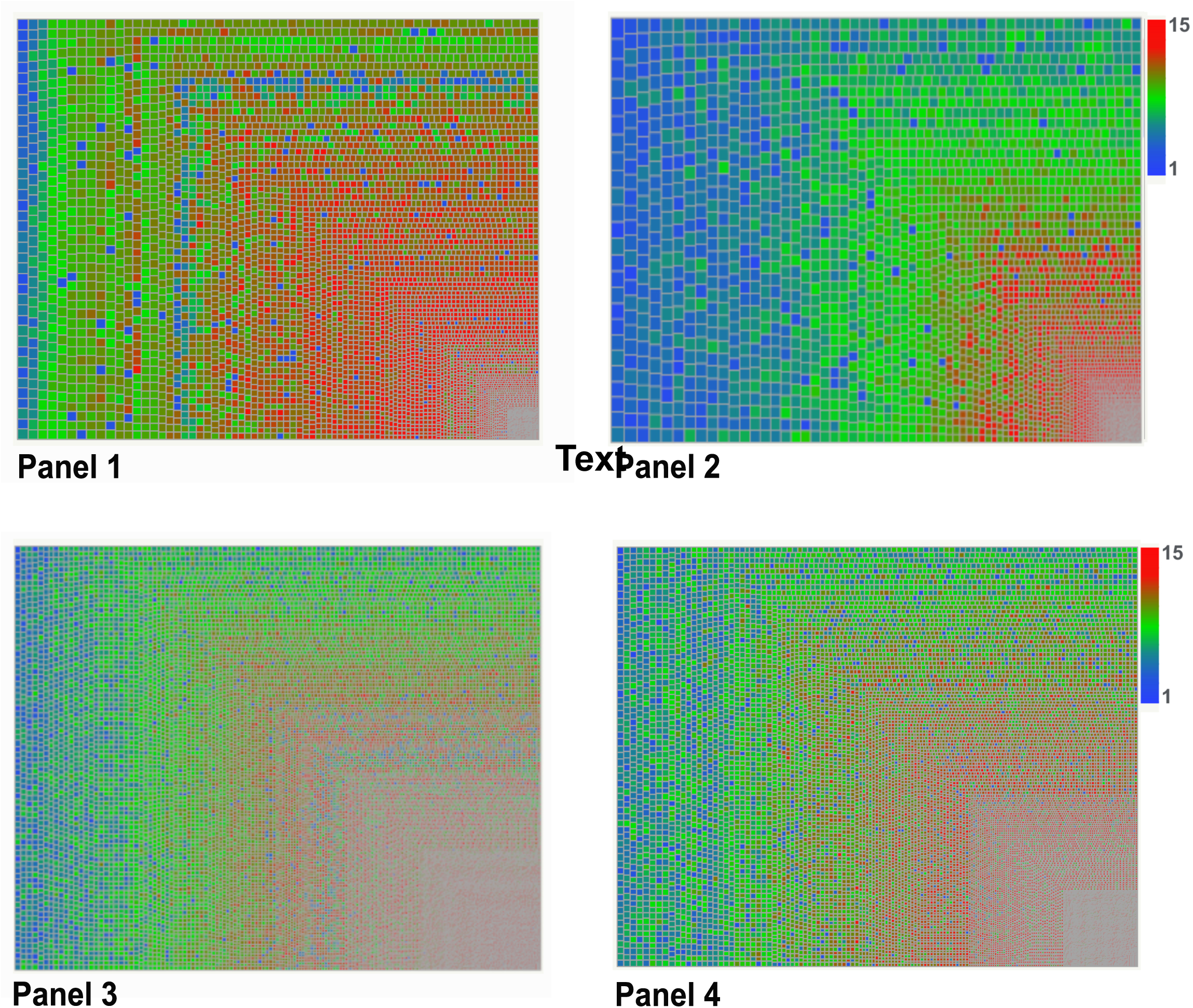
Mosaic plots for each antibody panel, in which the frequency of each immunophenotyped defined by combinatorics is depicted by the size of its tile. The immunophenotypes are arranged such that the most frequent populations are in the upper left corner of the graphic, while the least frequent are in the lower right corner. The color scale denotes the number of markers used to describe each immunophenotype.

**Supplemental Figure 3.**
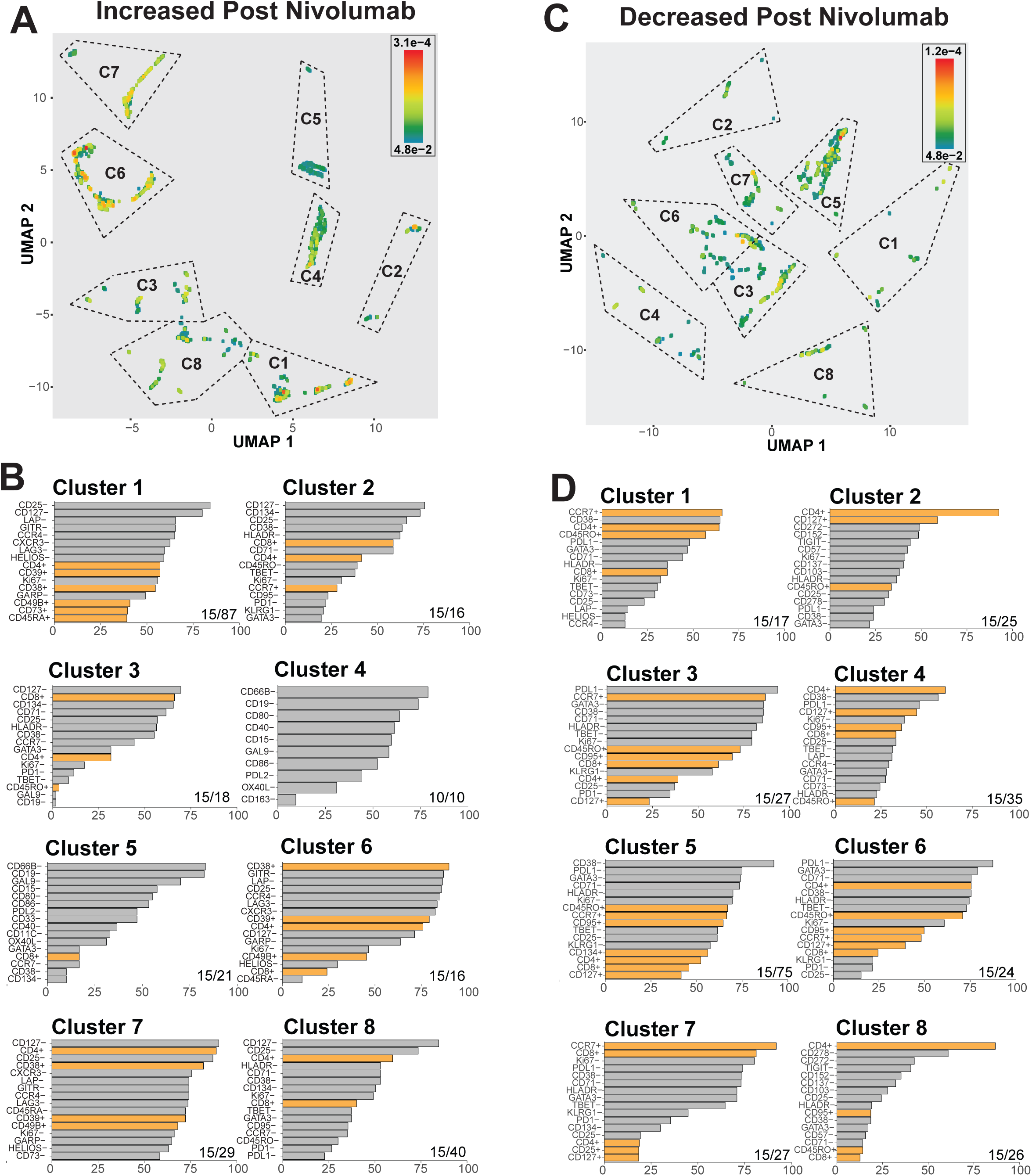
Peripheral immunophenotypes that change after nivolumab. **(A)** Immunophenotypes significantly increased post nivolumab (p<0.05, Wilcoxon signed-rank test) were reduced to a two-dimensional graph by Uniform Manifold Approximation and Projection (UMAP) by using frequency values for all patient samples as n-dimension variables. Clusters were determined by K-Means. Each dot represents a significant immunophenotype and is colored by the p-value. **(B)** The frequency for the top markers appearing in each cluster are graphed. Positive markers (e.g. CD38+) are shown in orange bars and negative markers (e.g. CD38-) are shown in grey bards. The total number of markers appearing is shown in the bottom right of each panel. **(C, D)** Immunophenotypes significantly decreased post nivolumab are likewise graphed and the cluster phenotypes shown.

**Supplemental Figure 4.**
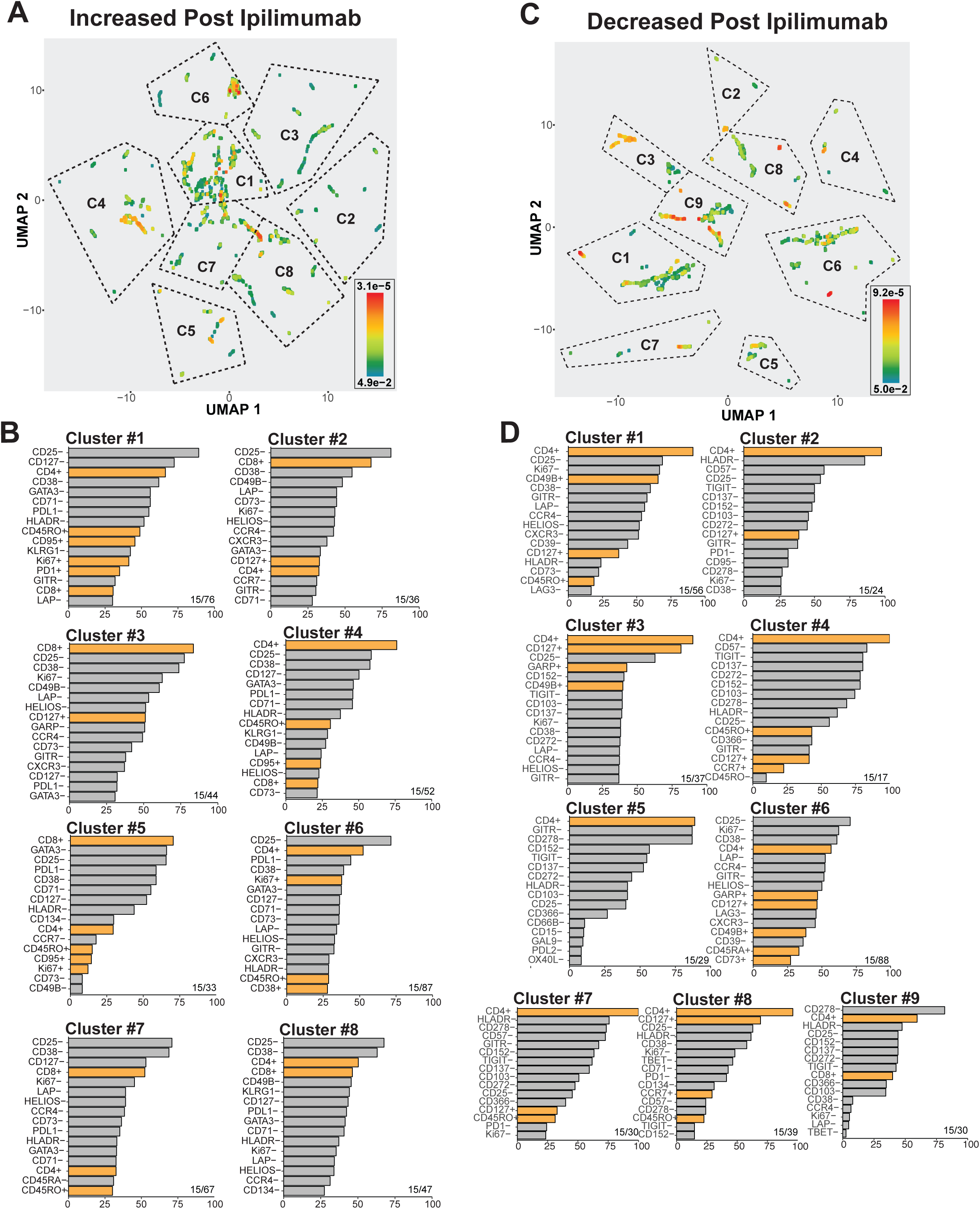
Peripheral immunophenotypes that change after ipilimumab. **(A)** Immunophenotypes significantly increased post ipilimumab (p<0.05, Wilcoxon signed-rank test) were reduced to a two-dimensional graph by UMAP by using frequency values for all patient samples as n-dimension variables. Clusters were determined by K-Means. Each dot represents a significant immunophenotype and is colored by the p-value. **(B)** The frequency for the top markers appearing in each cluster are graphed. Positive markers are shown in orange bars and negative markers are shown in grey bards. The total number of markers appearing is shown in the bottom right of each panel. **(C, D)** Immunophenotypes significantly decreased post ipilimumab are likewise graphed and the cluster phenotypes shown.

**Supplemental Figure 5.**
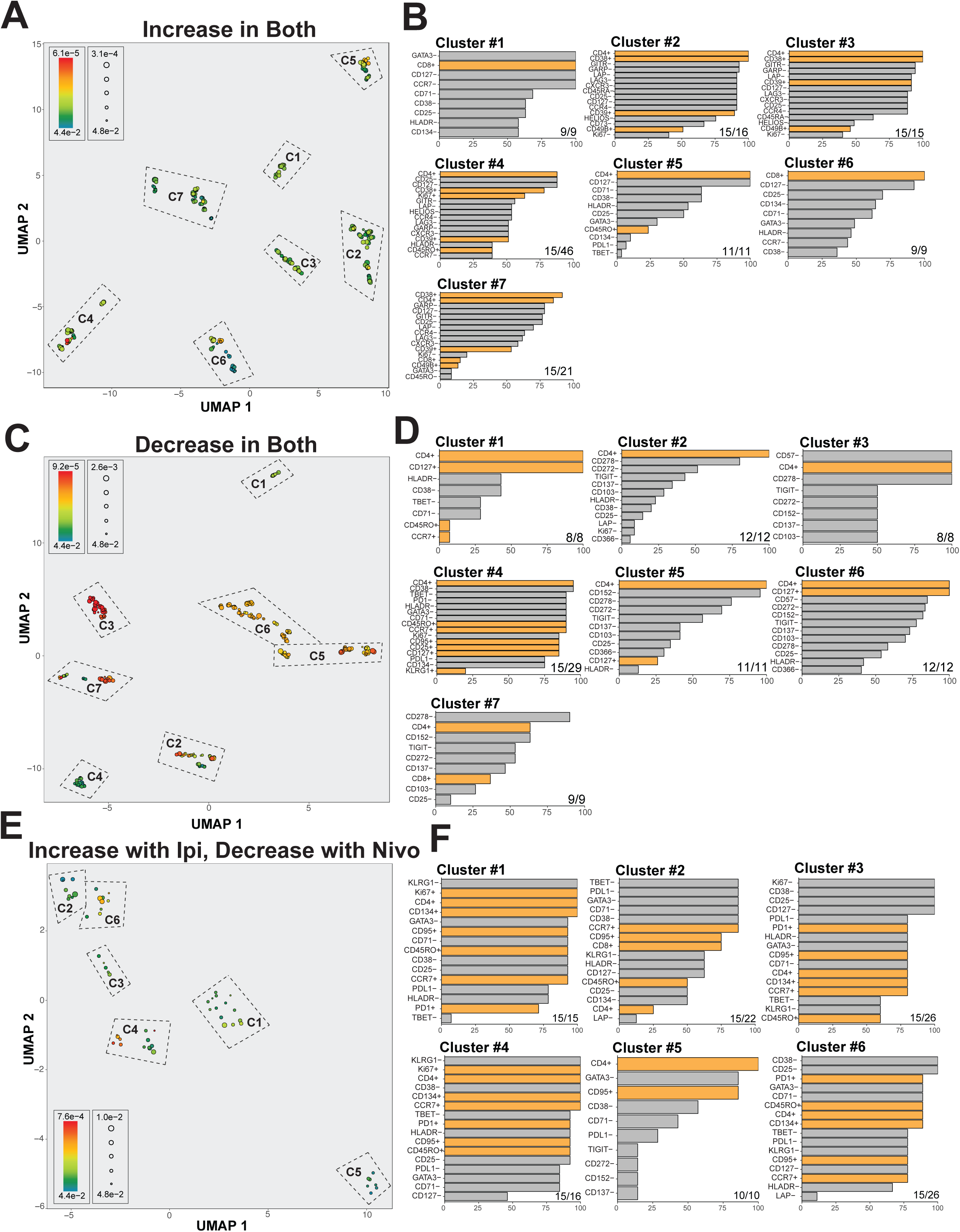
Overlap in nivolumab- and ipilimumab-induced peripheral immunophenotypic changes. **(A)** The 281 immunophenotypes found to be significantly increasing both post ipilimumab and post nivolumab (p<0.05, Wilcoxon signed-rank test) were reduced to a two-dimensional graph by UMAP by using frequency values for all patient samples as n-dimension variables. Clusters were determined by K-Means. Each dot represents a significant immunophenotype and is colored by the p-value of change after nivolumab treatment and is sized by the p-value of change after ipilimumab treatment. **(B)** The frequency for the top markers appearing in each cluster are graphed. Positive markers are shown in orange bars and negative markers are shown in grey bards. The total number of markers appearing is shown in the bottom right of each panel. **(C, D)** The 244 immunophenotypes significantly decreased both post nivolumab and post ipilimumab are likewise graphed and the cluster phenotypes shown. **(E, F)** The 56 immunophenotypes significantly decreased both post nivolumab and increased post ipilimumab are likewise graphed and the cluster phenotypes shown.

**Supplemental Figure 6.**
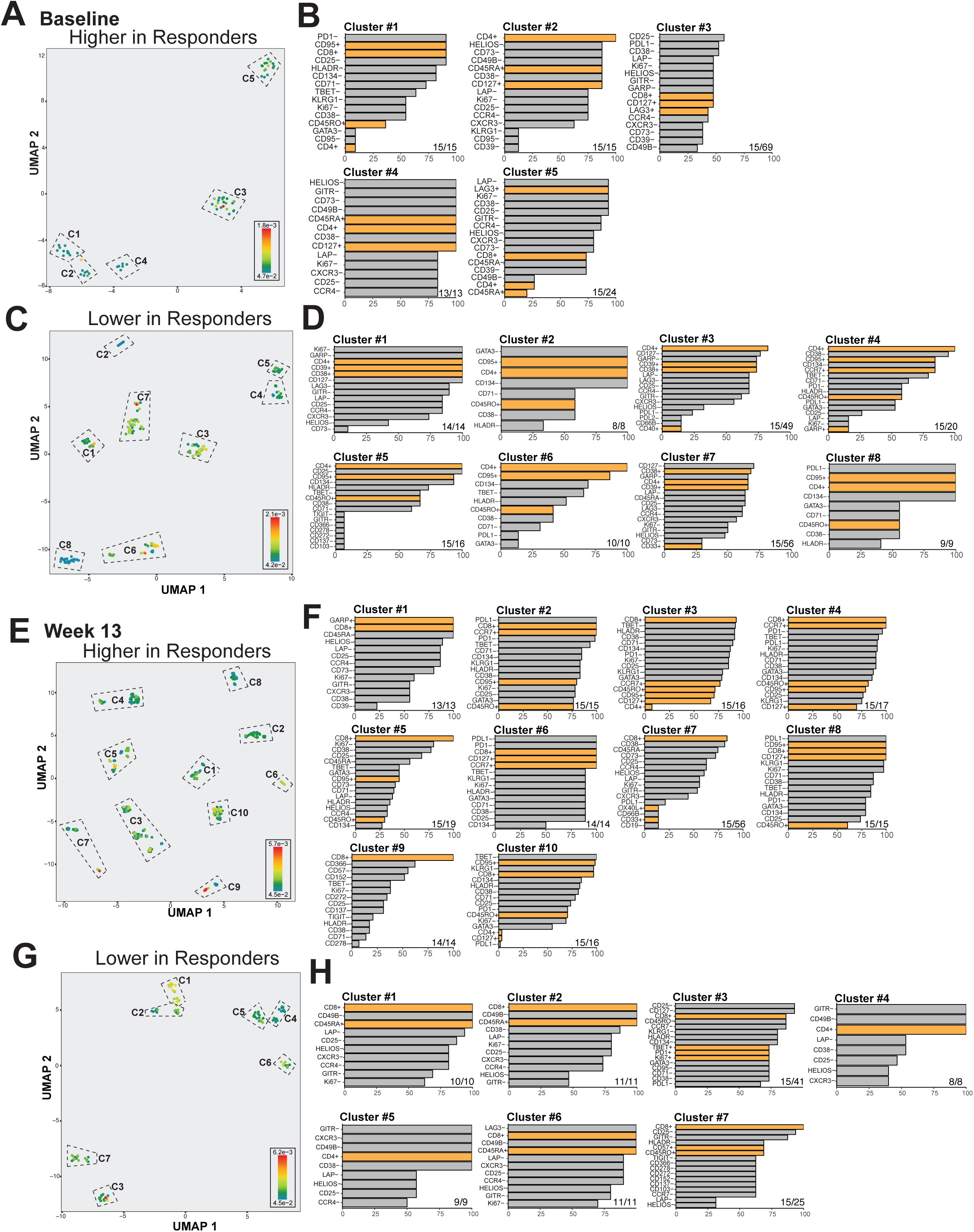
Biomarkers associated with patient response and survival after sequential NIVO>IPI therapy. **(A)** Baseline immunophenotypes significantly higher in nivolumab>ipilimumab responding patients relative to progressing patients were reduced to a two-dimensional graph by UMAP by using frequency values for all patient samples as n-dimension variables. Clusters were determined by K-Means. Each dot represents an immunophenotype and is colored by p-value. **(B)** The frequency for the top markers appearing in each cluster are graphed. Positive markers are shown in orange bars and negative markers are shown in grey bards. The total number of markers appearing is shown in the bottom right of each panel. **(C, D)** Baseline immunophenotypes significantly lower in nivolumab>ipilimumab responding patients are likewise graphed and the cluster phenotypes shown. **(E,F)** Post-nivolumab immunophenotypes significantly higher in nivolumab>ipilimumab responding patients are likewise graphed and the cluster phenotypes shown. **(G, H)** Post-nivolumab immunophenotypes significantly lower in NIVO>IPI responding patients are likewise graphed and the cluster phenotypes shown.

**Supplemental Figure 7.**
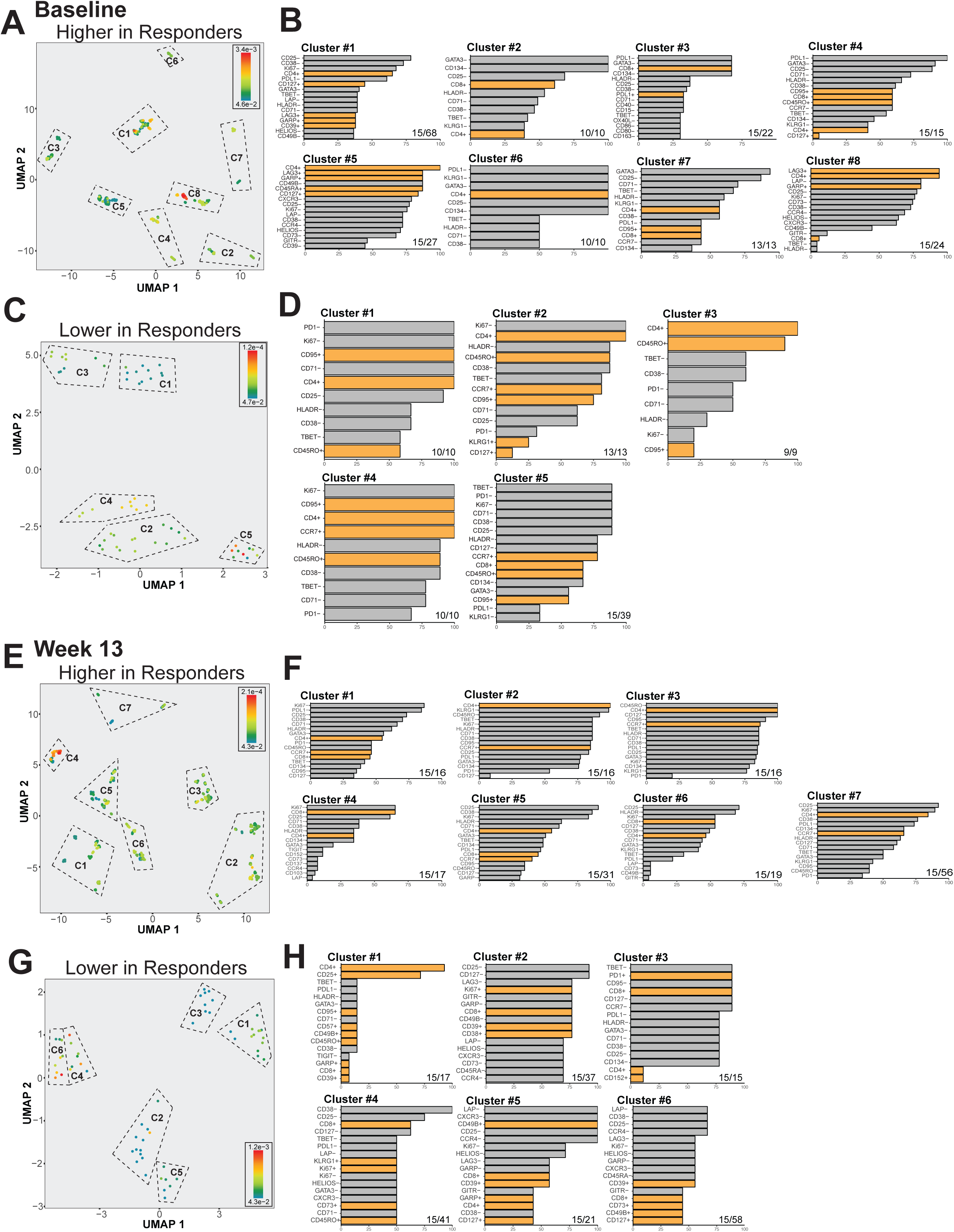
Biomarkers associated with patient response and survival after sequential IPI>NIVO therapy. **(A)** Baseline immunophenotypes significantly higher in ipilimumab>nivolumab responding patients relative to progressing patients were reduced to a two-dimensional graph by UMAP by using frequency values for all patient samples as n-dimension variables. Clusters were determined by K-Means. Each dot represents an immunophenotype and is colored by p-value. **(B)** The frequency for the top markers appearing in each cluster are graphed. Positive markers are shown in orange bars and negative markers are shown in grey bards. The total number of markers appearing is shown in the bottom right of each panel. **(C, D)** Baseline immunophenotypes significantly lower in ipilimumab-nivolumab responding patients are likewise graphed and the cluster phenotypes shown. **(E, F)** Post-ipilimumab immunophenotypes significantly higher in ipilimumab>nivolumab responding patients are likewise graphed and the cluster phenotypes shown. **(G, H)** Post-ipilimumab immunophenotypes significantly lower in IPI>NIVO responding patients are likewise graphed and the cluster phenotypes shown.

**Supplemental Figure 8.**
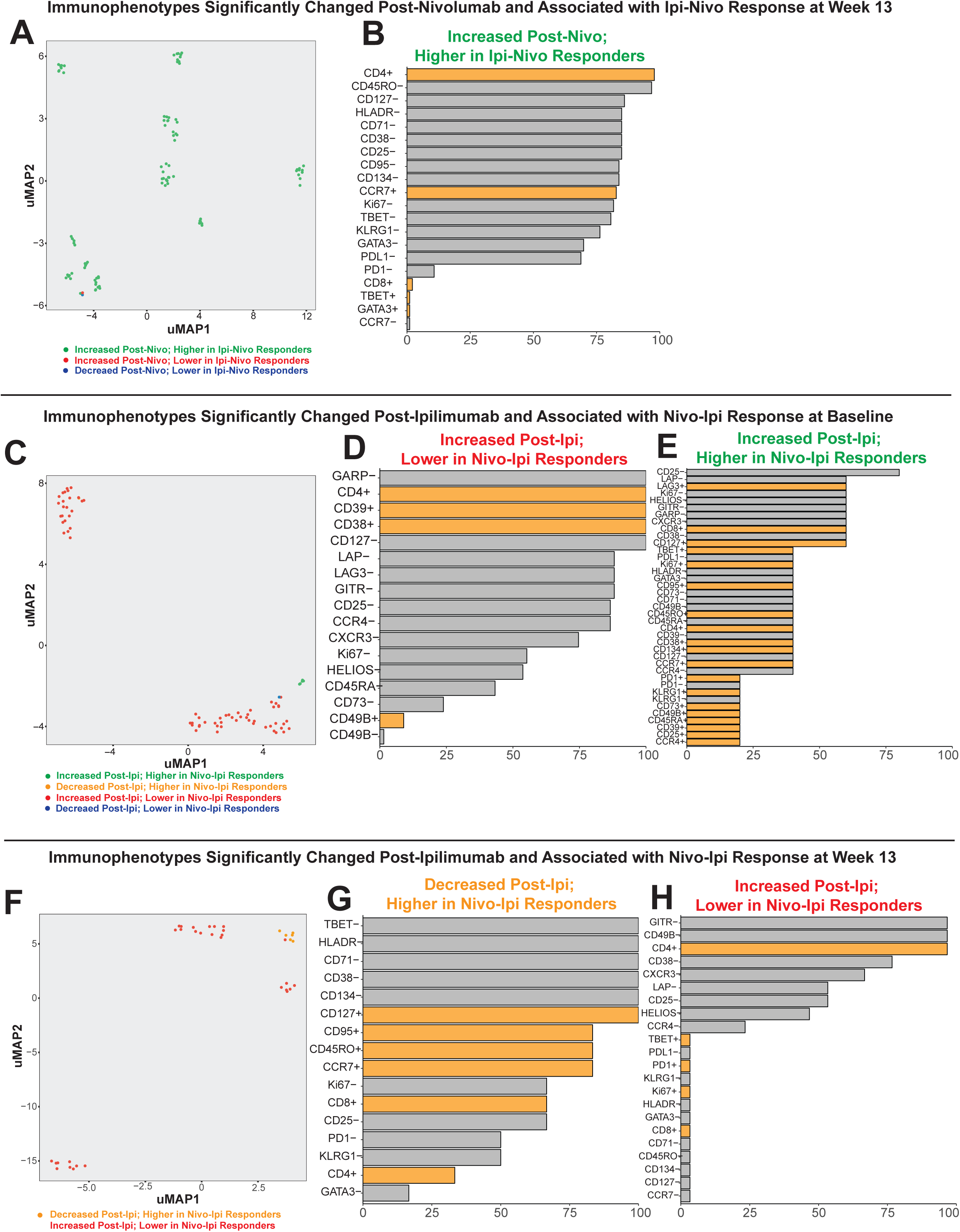
Immunophenotypes that are altered by Nivolumab or ipilimumab and overlap with biomarkers of patient outcomes. **(A)** Immunophenotypes significantly changed post-nivolumab and associated with response/survival in IPI>NIVO treated patients at week 13 were reduced to a two-dimensional graph by UMAP by using frequency values for all patient samples as n-dimension variables. Clusters were determined by K-Means. Each dot represents an immunophenotype and is colored by directions of change after nivolumab and relative representation in IPI>NIVO responding patients. Immunophenotypes increasing post-nivolumab and higher in IPI>NIVO responding patients are shown in green. Immunophenotypes increasing post-nivolumab and lower in IPI>NIVO responding patients are shown in red. Immunophenotypes decreasing post-nivolumab and lower in IPI>NIVO responding patients are shown in green. **(B)** The frequency for all markers appearing in immunophenotypes increasing post-nivolumab and higher in IPI>NIVO responding patients are plotted. **(C)** Immunophenotypes significantly changed post-ipilimumab and associated with response/survival in NIVO>IPI treated patients at baseline are likewise plotted. **(D)** The frequency for all markers appearing in immunophenotypes increasing post-ipilimumab and lower in NIVO>IPI responding patients are plotted. **(E)** The frequency for all markers appearing in immunophenotypes increasing post-ipilimumab and higher in NIVO>IPI responding patients are plotted. **(F)** Immunophenotypes significantly changed post-ipilimumab and associated with response/survival in NIVO>IPI treated patients at week 13 are plotted as in A and C. **(G)** The frequency for all markers appearing in immunophenotypes decreased post-ipilimumab and higher in NIVO>IPI responding patients are plotted. **(H)** The frequency for all markers appearing in immunophenotypes increasing post-ipilimumab and lower in NIVO>IPI responding patients are plotted.

**Supplemental Table 1.**
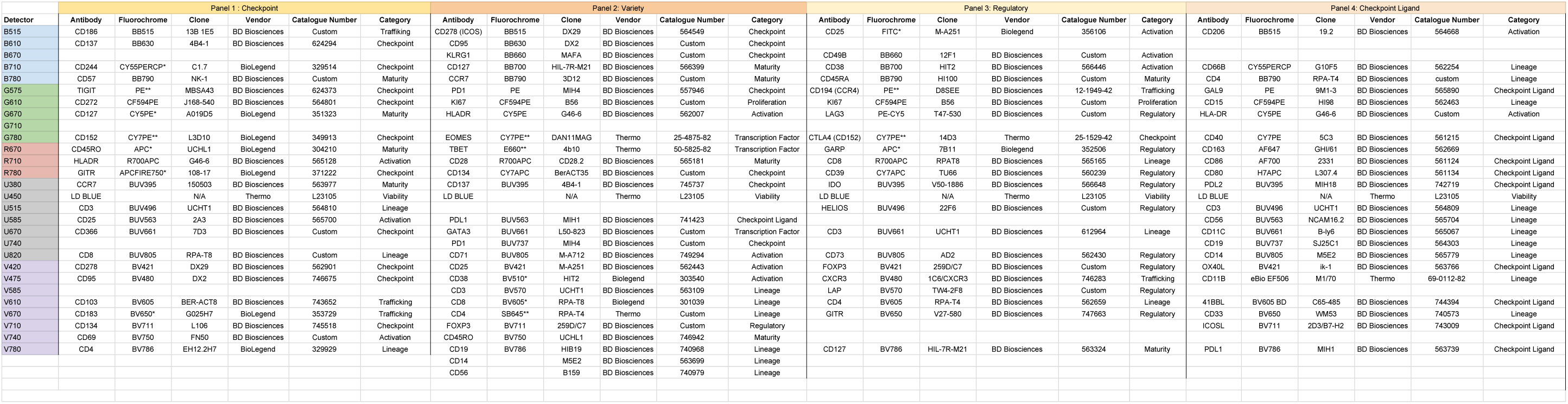
Flow cytometry staining panels.

